# SCF^FBXW7^ regulates G2-M progression through control of CCNL1 ubiquitination

**DOI:** 10.1101/2022.09.26.509608

**Authors:** Siobhan O’Brien, Susan Kelso, Zachary Steinhart, Stephen Orlicky, Monika Mis, Yunhye Kim, Sichun Lin, Frank Sicheri, Stephane Angers

## Abstract

*FBXW7*, which encodes a substrate specific receptor of an SCF E3 ligase complex, is a frequently mutated human tumor suppressor gene known to regulate the post-translational stability of various proteins involved in cellular proliferation. Here, using genome-wide CRISPR screens we report a novel synthetic lethal genetic interaction between *FBXW7* and *CCNL1* and describe CCNL1 as a new substrate of the SCF-FBXW7 E3 ligase. Further analysis showed that the CCNL1-CDK11 complex is critical at the G2-M phase of the cell cycle since defective CCNL1 accumulation, resulting from *FBXW7* mutation, leads to shorter mitotic time. Cells harboring *FBXW7* loss-of-function mutations are hypersensitive to treatment with a CDK11 inhibitor, highlighting a genetic vulnerability that could be leveraged for cancer treatment.

## Introduction

F-box and WD repeat domain containing protein 7 (FBXW7) is a substrate specific recognition module of a Skp1-Cul1-Fbox (SCF) E3 ligase complex named SCF^FBXW7^ (Welcker and Clurman, 2008). SCF E3 ligases target several proto-oncogenes for ubiquitin-dependent proteasomal degradation to regulate cell cycle progression and proliferation (Koepp, 2001; Reed et al., 2004; Wei et al., 2005; Welcker et al., 2004b; Yeh et al., 2018). One of the better characterized substrates of SCF^FBXW7^ is cyclin E, a master regulator of the G1-S cell cycle phase transition (Koepp, 2001; Ohtsubo et al., 1995). Loss of *FBXW7* and concomitant accumulation of cyclin E deregulates the cell cycle (Reed et al., 2004), enhances DNA replication (Minella et al., 2008), and causes genomic instability across many different cell types (Loeb et al., 2005; Minella et al., 2007; Rajagopalan et al., 2004; Spruck et al., 1999). c-MYC is another well-studied substrate of SCF^FBXW7^ (Welcker et al., 2004a, 2004b). Although not directly implicated in regulation of cell-cycle phase transition, c-MYC inhibits many inhibitors of the cell cycle including p21 and p27 which negatively regulate the G1-S transition checkpoint (Mateyak et al., 1999; Perez-Roger et al., 1999).

Given its fundamental roles in cell cycle control, *FBXW7* inactivating mutations that lead to substrate stabilization are common across a wide-range of cancers, with highest mutational frequency in uterine, cervical and intestinal cancers (Yeh et al., 2018). The most common mutations are within *FBXW7* exons encoding WD-repeats, which function as the substrate-recognition domain. These mutations lead to defective post-translational control of proto-oncogene abundance and hence promote cancer progression. Biochemical studies have confirmed that the WD-repeat hotspot mutations (R465C/H/L, R479P/Q/*, and R505C/G/H) are loss of function mutations that disrupt substrate binding (Orlicky et al., 2003; Tang et al., 2007). In addition to cyclin E and c-Myc, SCF^FBXW7^ has also been implicated in the control of other potentially oncogenic substrates, including c-Jun (Nateri et al., 2004; Wei et al., 2005), and Notch1 (Gupta-Rossi et al., 2001; Oberg et al., 2001; O’Neil et al., 2007; Tetzlaff et al., 2004).

Currently categorized as a transcriptional or non-canonical cyclin, CCNL1 (cyclin L1) was demonstrated to functionally regulate the spliceosome, along with its serine/threonine kinase partner cyclin-dependent kinase 11 (CDK11) (Chen et al., 2007, 2006; Loyer and Trembley, 2020). Consistent with the understanding of cyclin-CDK biology, the primary role of CCNL1 is to promote CDK11 activity (Loyer and Trembley, 2020). CCNL1 has been proposed as a candidate oncogene in head and neck cancer due to high levels of chromosomal amplification that correlates with poor overall survival in patients (Muller et al., 2006; Redon et al., 2002; Sticht et al., 2005). Amplification of *CCNL1* is also associated with poor prognosis in uterine cancer (Mitra et al., 2010). CDK11 has several isoforms, with the p58 version generated via usage of an internal ribosomal entry site within CDK11 mRNA transcripts at the G2/M cell cycle transition (Cornelis et al., 2000). Recently, the CCNL1-CDK11 complex has been implicated in cytokinesis, with CDK11-p58 kinase activity required for abscission, the final stage of mitosis (Renshaw et al., 2019). Despite a poor understanding of the role of CCNL1 in cancer initiation and progression, several studies have established CDK11 as an important regulator of cancer cell proliferation and that its loss of function is lethal in many cancer types (Ahmed et al., 2019; Liu et al., 2016). A novel selective CDK11 inhibitor, OTS964, was recently serendipitously identified (Lin et al., 2019) and offers a potentially tractable therapeutic opportunity for cancers, perhaps in contexts where tumor cells are reliant on CDK11 activity.

In the present study, we combine drug selection with gene-editing of pancreatic cancer cells to rewire their growth dependency in such a way to be exquisitely reliant on the loss of FBXW7 activity. This engineered model was required to ensure that cell growth was dependent on this single *FBXW7* mutation in order to hone in the molecular mechanisms underlying this frequent cancer causing alteration. Using genome-wide CRISPR fitness screens performed in isogenic cell lines, we uncovered a novel synthetic lethal interaction between *FBXW7* and *CCNL1*. Given that SCF^FBXW7^ is known to control the levels of other cyclins, we showed that CCNL1 is a novel substrate for this E3 ligase. Our findings suggest that the deregulation of this axis is frequent in human cancers and it culminates in the hyperactivation of CCNL1’s kinase partner CDK11, thereby uncovering a novel therapeutic opportunity.

## Results

### Loss of *FBXW7* induces resistance to Wnt-inhibition in a Wnt-addicted cell line

A subset of pancreatic adenocarcinoma harbor inactivating mutations within *RNF43*, a negative regulator of the Wnt-ßcatenin pathway (Jiang et al., 2013). As a result of *RNF43* mutation cells express high levels of Frizzled receptors and are exquisitely dependent on autocrine Wnt-ßcatenin signaling for growth as highlighted by their hypersensitivity to LGK974, a small molecule inhibitor of porcupine (PORCN) that blocks secretion and function of Wnt proteins (**Figure 1A,B**), as well as to anti-Frizzled blocking antibodies (Steinhart et al., 2017). To pre-emptively study mechanisms of resistance to Wnt inhibitors, we conducted a genome-wide CRISPR suppressor screen, in the *RNF43* mutant cell line HPAF-II. This experiment identified gene knockouts that overcome the growth arrest phenotype induced by LGK974 (**Figure 1C,D**). Predictably, the screen identified the well-known negative regulators of ßcatenin signaling, *AXIN1, APC*, and *CSNK1A1* (**Figure 1D**) as these mutations all lead to ligand-independent ßcatenin stabilization and regulation of gene expression and hence bypass the requirement for autocrine Wnt ligands (Amit et al., 2002; Liu et al., 2002; Steinhart et al., 2017; Su et al., 2008). The genes above represented nearly 20% of the total amount of next generation sequencing reads in the suppressor screen highlighting their strong negative regulatory functions during Wnt-ßcatenin signaling. In addition to these genes, the screen also uncovered tumor suppressor F-box protein *FBXW7*, which functions as a substrate recognition subunit within a Skp1-Cullin-Fbox (SCF) E3 ligase complex and is a well-studied tumor suppressor gene (Mao et al., 2004; Yeh et al., 2018). *APC* and *FBXW7* HPAF-II knockout cells were generated using CRISPR and selected using selective pressure with LGK974 (**Figure 1E, EV1A,B**). Strikingly, whereas *FBXW7* knockout cells were confirmed to be resistant to LGK974-mediated cell cycle arrest (**Figure 1F**), with relative confluence reduced to 9.1% in the wild-type, 28% in *FBXW7*^-/-^ and unchanged in *APC*^-/-^ in the presence of LGK974 **(Figure 1F)**, only *APC*^-/-^ cells maintained high ßcatenin levels in the presence of LGK974 (**Figure 1G**). This data confirms that in HPAF-II cells, FBXW7 itself is not regulating ßcatenin levels as was reported in a different context (Jiang et al., 2016). We conclude that *FBXW7*^-/-^ cells are partially resistant to *PORCN* inhibitor-mediated growth arrest by a mechanism that does not involve reactivation of downstream ßcatenin signaling.

**Figure 1.**
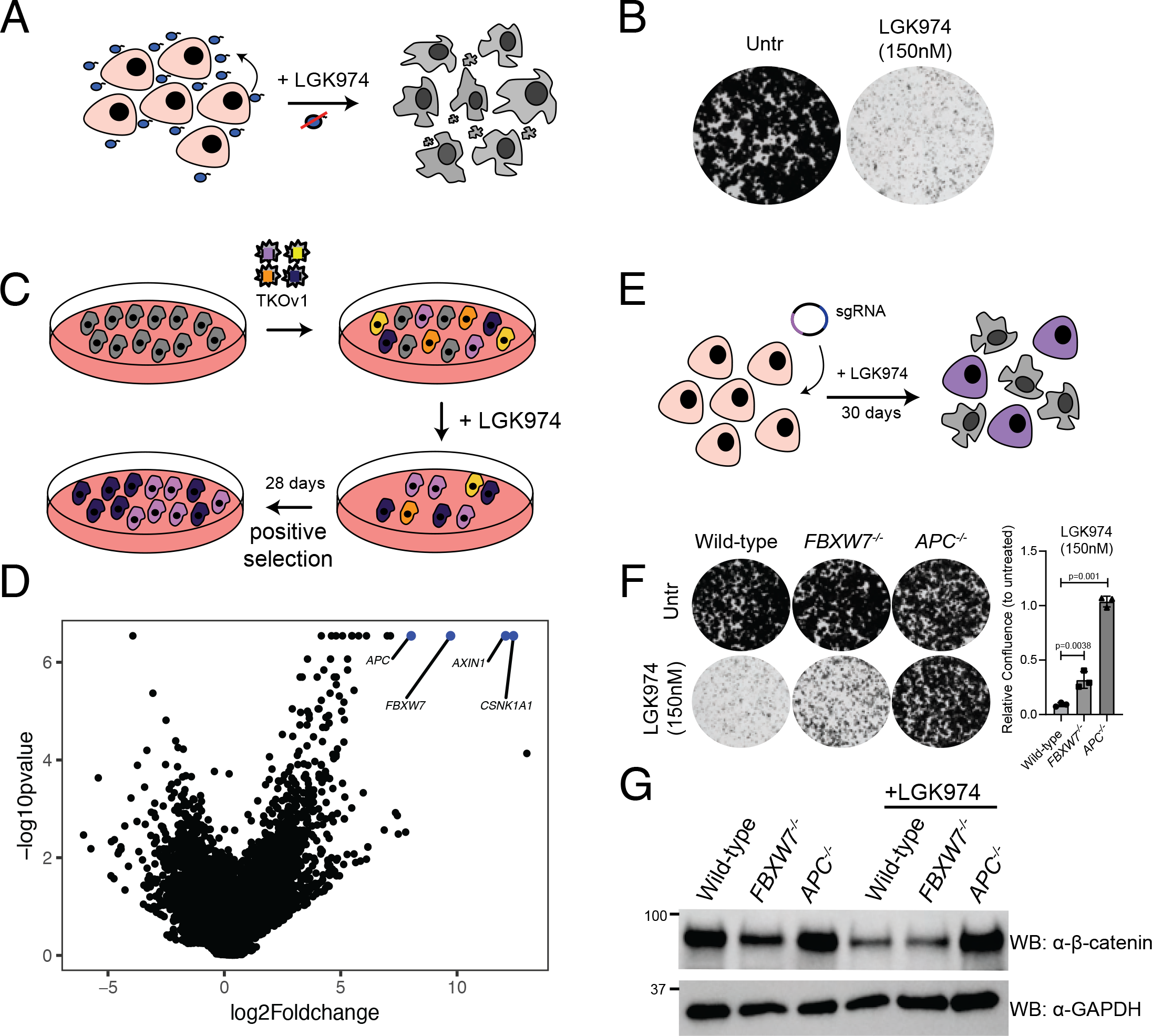
*FBXW7* loss-of-function bypasses requirement for autocrine Wnt signaling in Wnt-addicted PDAC cells. A HPAF-II cells contain an autocrine Wnt signaling loop, where Wnt secreted by cells is required for the cells to grow. Treatment with the Porcupine (PORCN) inhibitor LGK974 (which inhibits Wnt secretion) leads to cell cycle arrest. B Clonogenic growth assay in control HPAF-II cells or in cells treated with 150nM LGK974 for 10 days; representative of three independent replicates. C Schematic representation of the positive selection screen conducted to identify genes involved in LGK974 sensitivity. D Volcano plot with results of the positive selection screen identifying *FBXW7* along with *AXIN1, CSNK1A1*, and *APC* as genes overcoming LGK974-induced cell cycle arrest when knocked out. E Schematic representation of *FBXW7*^*-/-*^ and *APC*^*-/-*^ cell line generation in HPAF-II using LGK974 treatment to enrich edited cells. F Clonogenic growth assay of indicated HPAF-II cells left untreated or in the presence of 150nM LGK974 for 10 days; representative of three independent replicates. Quantification of n=3 biological replicates, one-way ANOVA. G Immunoblot of cytoplasmic βcatenin expression from lysates of HPAF-II cells from indicated genotype, treated with vehicle or 100nM LGK974 for 48h; representative of three biological replicates.

### *CCNL1* loss-of-function is synthetic lethal with *FBXW7* mutation

To reveal the growth mechanisms dysregulated in *FBXW7*^-/-^ cells that underlie the resistance to LGK974 treatment, we performed isogenic CRISPR fitness screens in wild-type and *FBXW7*^-/-^ HPAF-II cells (**Figure 2A**). We then used the BAGEL algorithm to calculate a Bayes factor (BF) for each gene (Hart and Moffat, 2016). BF is a confidence score that knockout of a specific gene causes a decrease in fitness where high BF indicates increased confidence that the knockout of the gene results in a decrease in fitness. We then derived a differential fitness score for each gene by subtracting the BF scores obtained in *FBXW7*^-/-^ and wild-type cells and plotted the differential Z-score (**Figure 2B, EV1C,D**). Confirming that *FBXW7*^-/-^ cells have evaded a requirement for Wnt-ßcatenin signaling, several of the genes we previously identified as fitness genes in HPAF-II cells (*FZD5, PORCN, TCF7L2, WLS, CTNNB1*) (Steinhart et al., 2017) have negative Z-scores, indicating a greater fitness defect observed in wild-type cells. Interestingly, the gene with the highest positive differential Z-score was the poorly characterized cyclin family member cyclin L1 (*CCNL1*) (**Figure 2B** and individual CCNL1 gRNA dropouts comparison in **Figure 2C**). To validate these results, we performed multicolor cell competition assays with HPAF-II or HPAF-II *FBXW7*^-/-^ mutant cells expressing a control gRNA targeting *AAVS1* (labeled with mCherry) or two independent gRNAs targeting *CCNL1* (labeled with GFP) and showed that mCherry cells outcompeted GFP cells at a much faster rate in the absence of *FBXW7*, with a 50% reduction in GFP-expressing cells by day 4, with wild-type cells largely unaffected until day 16 (**Figure 2D, EV1E**). Supporting these results, when infected with lentivirus encoding for Cas9 and gRNAs targeting *CCNL1*, HPAF-II *FBXW7*^-/-^ mutant cells proliferated at a much slower rate when compared to wild-type HPAF-II cells (**Figure 2E, EV1E**). We conclude that loss of *CCNL1* is synthetic lethal with *FBXW7* mutation.

**Figure 2.**
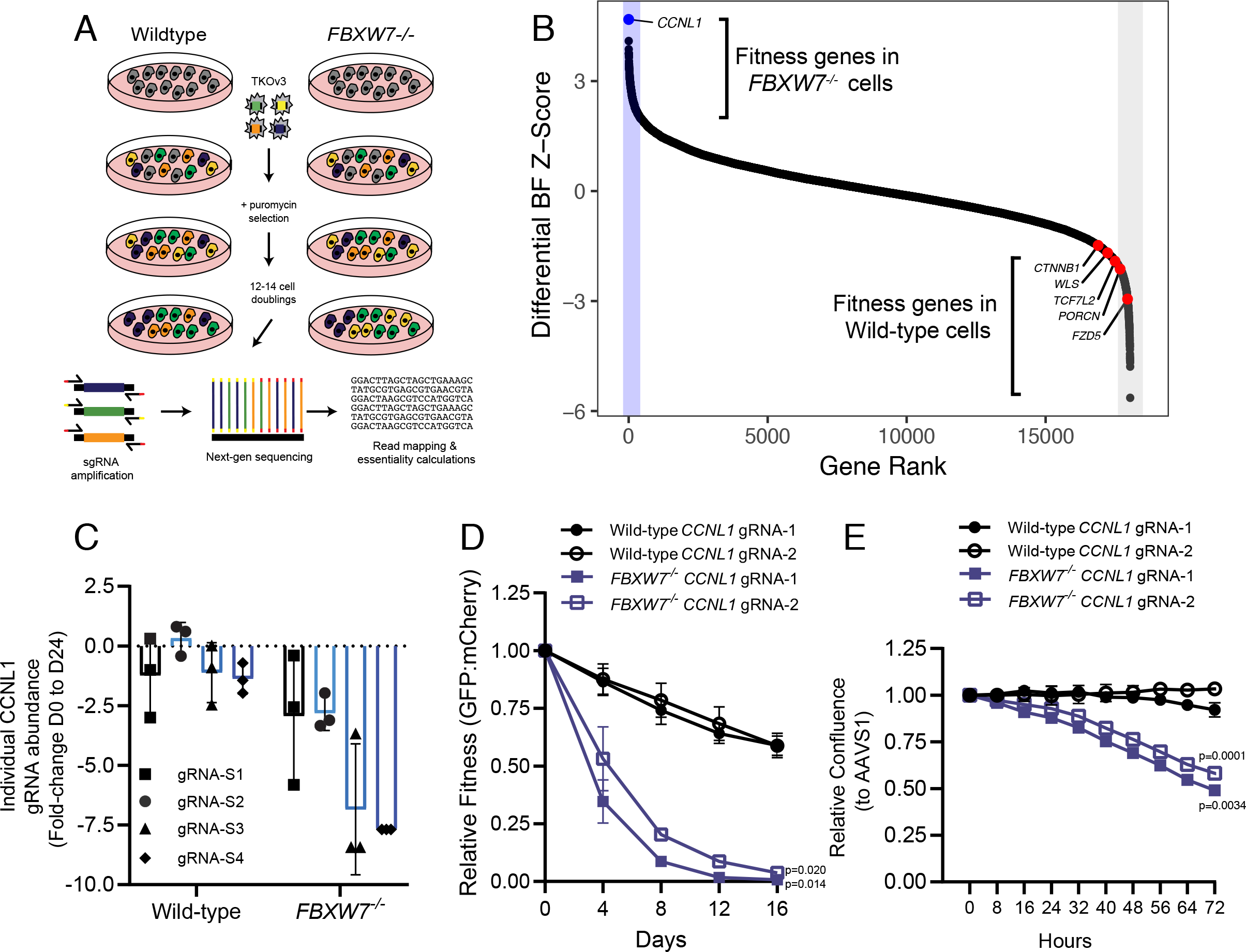
Genome-wide CRISPR screen performed in isogenic wild-type and *FBXW7*^*-/-*^ HPAF-II cells identifies a synthetic lethal genetic interaction. A Schematic representation of genome-wide CRISPR-Cas9 dropout screens performed in isogenic wild-type and *FBXW7*^*-/-*^ HPAF-II cell lines. B Differential Bayes Factor Z-score plot comparing wild-type and *FBXW7*^*-/-*^ genome-wide dropout screens. C Fold-change abundance of individual sgRNA targeting *CCNL1* during the genome-wide dropout screens from day 0 to day 24, n=3 technical replicates per sgRNA, mean ± SEM. D Multicolour-competition assay in both wild-type and *FBXW7*^*-/-*^ cell lines, using mCherry- DAVS1 and GFP-GOI, normalized to AAVS1 control cells at each time-point (n=3 independent replicates), mean ± SEM, one-way ANOVA. E Proliferation assays in wildtype and *FBXW7*^*-/-*^ cell lines show that knockout of CCNL1 preferentially affects *FBXW7*^*-/-*^ cells, normalized to AAVS1 control (n=3 independent replicates), mean ± SEM, one-way ANOVA

### CCNL1 is a substrate of the SCF^FBXW7^ E3 ligase

The well established role of SCF^FBXW7^ in regulating cyclin E stability hinted that one potential mechanism underlying the observed synthetic lethality phenotype is that CCNL1 is a novel substrate of the SCF^FBXW7^ E3 ligase. In support, we noted increased steady state CCNL1 levels in *FBXW7*^-/-^ cells when compared to wild-type cells (**Figure 3A, EV2A**). Cycloheximide chase further revealed that the half-life of CCNL1 was extended in *FBXW7*^-/-^ vs wild-type cells (**Figure 3B,C**). Expression of a dominant-negative Cul1 mutant (Van Rechem et al., 2011) induced stabilization of CCNL1 at steady state, further indicating that an SCF complex is involved in regulation of CCNL1 **(Figure 3D, EV2B)**. We next scanned the amino acid sequence of CCNL1 for the presence of a canonical Cdc4 phosphodegron (CPD) motif present in the majority of FBXW7 substrates (Nash et al., 2001; Orlicky et al., 2003) and identified a TPXXS sequence at position 325-329 (**Figure 3E**). Analysis of CPD peptide binding to purified FBXW7 using fluorescence polarization assays revealed a requirement for dual phosphorylation at both T325 and S329 within the CCNL1 CPD motif (**Figure 3F**), similar to what is seen for the FBXW7-Jun interaction (Wei et al., 2005), but in contrast to the FBXW7-cyclin E interaction where phosphorylation at the threonine residue is sufficient for maximal binding (Hao et al., 2007, and **Figure 3F**). Mutation of the TPXXS motif to VPXXA in the context of full length CCNL1 extended the half life of CCNL1 compared to wild-type proteins following cycloheximide chase (**Figure 3G,H**). Co-expression of HA-CCNL1 with FLAG-FBXW7 further reduced the half-life of CCNL1, while co-expression with the substrate binding mutant FBXW7^R465C^ fully stabilized CCNL1 expression **(Figure EV2G,H)**, demonstrating a direct role of FBXW7 in regulating CCNL1 expression.

**Figure 3.**
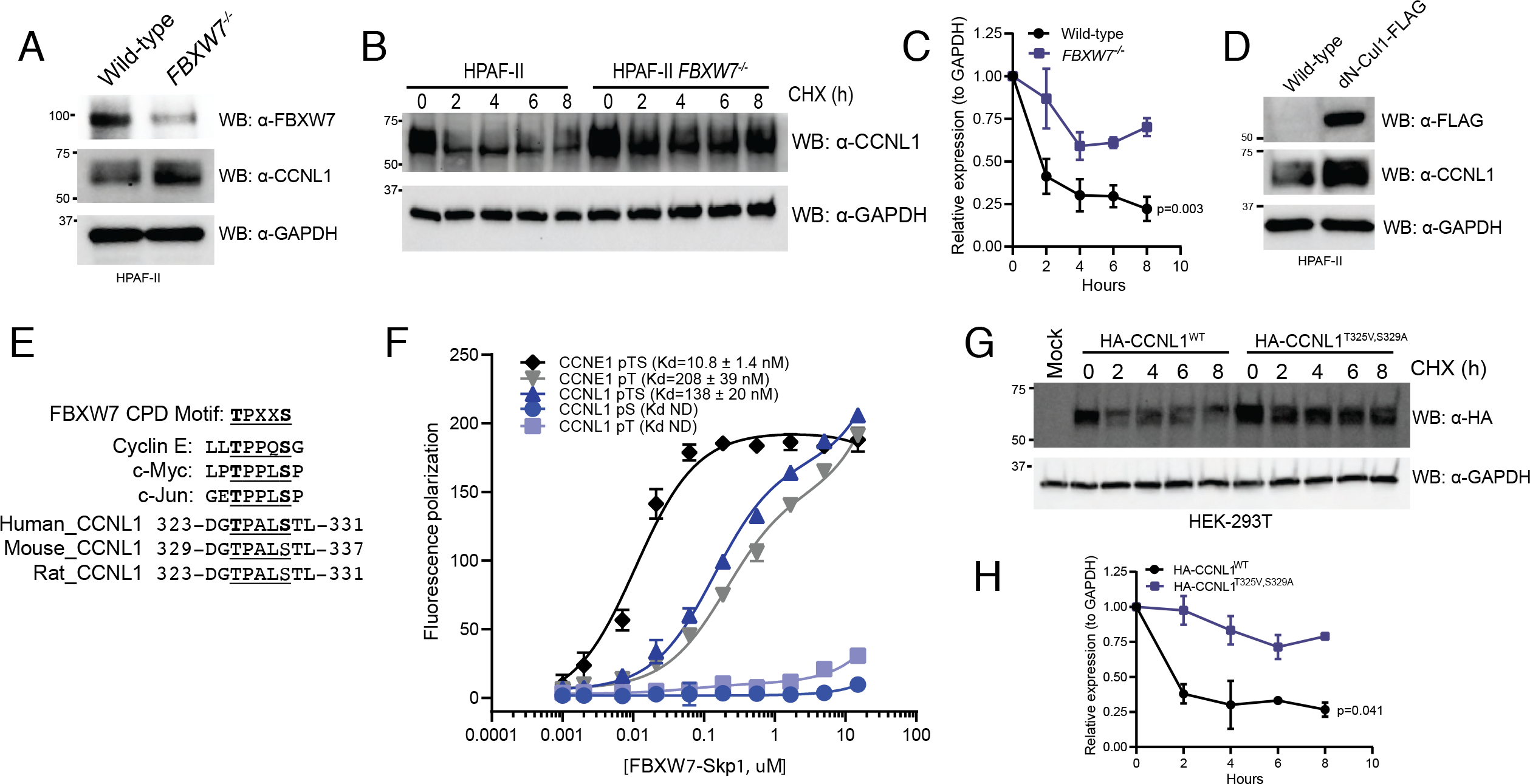
CCNL1 protein stability is mediated by SCF^FBXW7^. A Immunoblot of CCNL1 and FBXW7 expression in HPAF-II wild-type and *FBXW7*^*-/-*^ cells, representative blot of 3 independent replicates. B Immunoblot of CCNL1 expression following a cycloheximide chase in HPAF-II wild-type and *FBXW7*^*-/-*^ cells, representative blot of 3 independent replicates. C Quantification of cycloheximide chase in B, mean ± SEM of three independent replicates. Two-way ANOVA at T8. D Immunoblot of CCNL1 expression in HPAF-II wild-type and dN-Cul1 expressing cells, representative blot of 3 independent replicates. E Sequence alignment of well-characterized CPD degron motifs, and identification of potential FBXW7 phosphodegron motif in a conserved region of CCNL1. F Fluorescence polarization assay of CCNL1 and cyclin E peptides binding FBXW7-Skp1 complex, three independent replicates, mean ± SEM. G Immunoblot of lysates following cycloheximide treatment of HEK293T cells expressing wild-type or degron-mutated CCNL1, representative blot of 3 independent replicates. H Quantification of cycloheximide chase in G, mean ± SEM of three independent replicates. Two-way ANOVA at T8.

To verify that CCNL1 and FBXW7 are indeed interacting in cells, we performed an immunoprecipitation assay using overexpression of either FLAG-CCNL1 or FLAG-FBXW7, and detected interactions with endogenous FBXW7 and CCNL1 respectively **(Figure 4A,B)**. We next wished to perform *in vitro* ubiquitination assays but were faced with a roadblock in our multiple attempts to purify CCNL1 from either Sf9 or *E*.*coli* cultures. We therefore employed a cellular ubiquitination assay in HEK293T cells to confirm CCNL1 ubiquitination and the role of T325 and S329 in the process **(Figure 4C)**. A second cellular ubiquitination assay in HEK293T cells expressing a control sgRNA targeting *AAVS1*, or an sgRNA targeting *FBXW7* demonstrated that reducing FBXW7 expression reduced ubiquitination of CCNL1 **(Figure 4D, EV2C,D)**. Using HEK293T cells expressing an *FBXW7*-targeting gRNA, we added a gRNA-resistant FLAG-FBXW7 cDNA and assessed CCNL1 ubiquitination. The results indicated that re-expression of FBXW7 rescued CCNL1 ubiquitination **(Figure 4E)**. We detected an interaction between CCNL1 and endogenous CUL1 in HPAF-II cells, further supporting a role of the SCF complex in CCNL1 degradation. Interestingly, CUL4A was also co-immunoprecipitated with CCNL1 perhaps suggesting a role for this cullin in regulation of CCNL1 stability, similar to another FBXW7 substrate -Jun-which is degraded by both CUL1 and CUL4A based E3 ligases (Cang et al., 2007) **(Figure EV2E)**. We conclude that CCNL1 is a bona fide substrate of the SCF^FBXW7^ complex.

**Figure 4.**
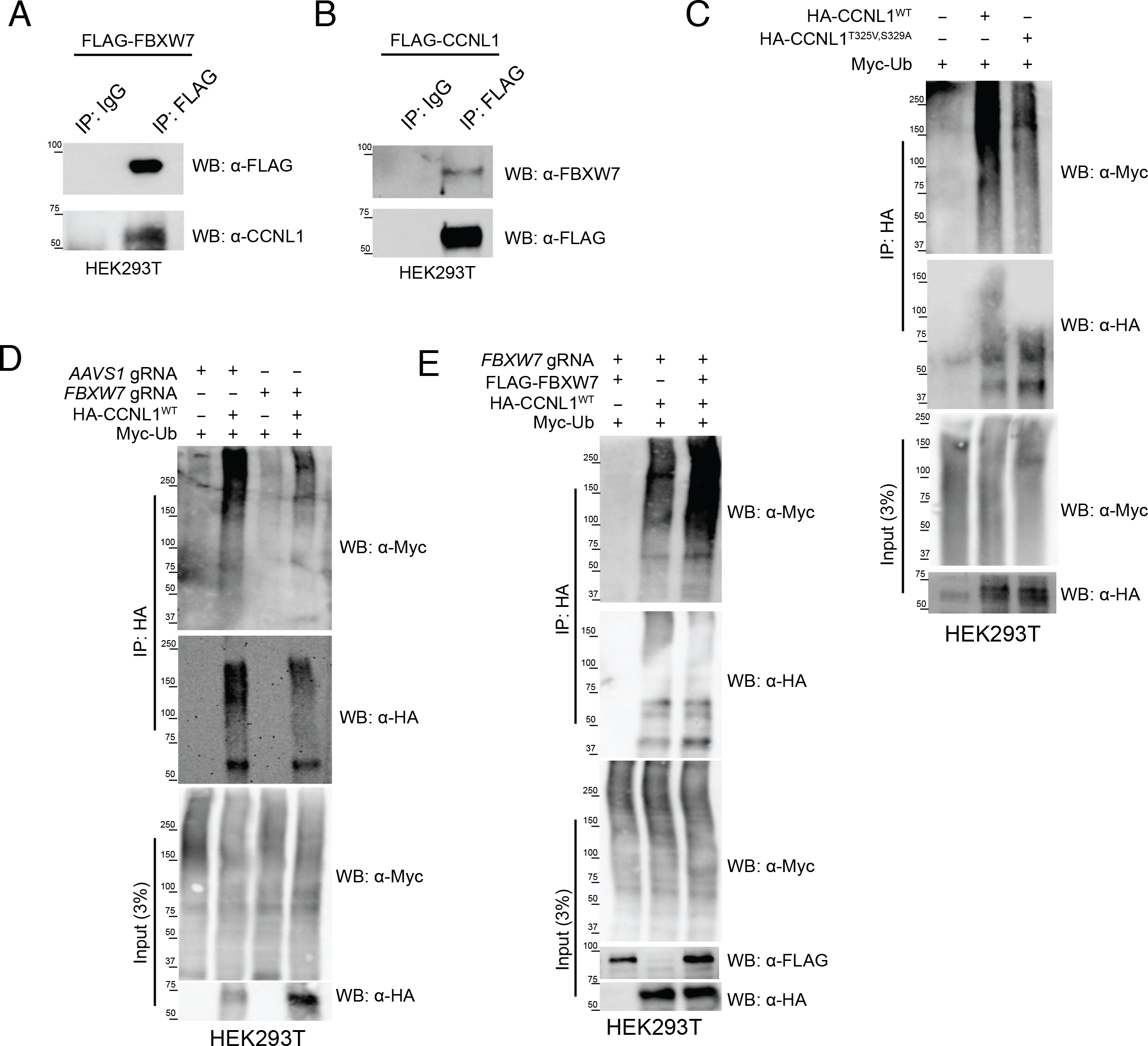
FBXW7 is involved in the ubiquitination of CCNL1. A Immunoblot of immunoprecipitation of FLAG-FBXW7 overexpressed in HEK293T cells, detecting endogenous CCNL1. Representative of three independent replicates. B Immunoblot of immunoprecipitation of FLAG-CCNL1 overexpressed in HEK293T cells, detecting endogenous FBXW7. Representative of three independent replicates. C Immunoblot of cellular ubiquitination assay demonstrating the requirement of the 325-TPALS-329 degron for ubiquitination of CCNL1. Representative of three independent replicates. D Immunoblot of cellular ubiquitination assay demonstrating the requirement of FBXW7 for ubiquitination of CCNL1. Representative of three independent replicates. E Immunoblot of cellular ubiquitination assay showing re-expression of FBXW7 in knockout cells leads to increased CCNL1 ubiquitination. Representative of three independent replicates.

### FBXW7 regulates G2-M progression through regulation of CCNL1 stability

To identify whether CCNL1 has a role in cell cycle progression like other SCF substrates, a cell cycle profile experiment was performed following release of cells that were first arrested in mitosis using nocodazole. In wild-type cells, CCNL1 expression oscillates in a pattern similar to cyclin B, supporting a potential role of CCNL1 in the G2-M phase of the cell cycle (**Figure 5A,B**). In contrast, in *FBXW7*^-/-^ cells the cycling of CCNL1 levels is lost, supporting the role of FBXW7 in targeting CCNL1 for degradation in a cell cycle dependent manner (**Figure 5A,B**). Considering the previously described role of CCNL1 and CDK11 in cytokinesis (Renshaw et al., 2019), we assessed whether upregulation of CCNL1 through either *FBXW7* loss-of-function or *CCNL1* overexpression affected normal progression through mitosis. First, to assess the cell cycle dynamics during logarithmic growth, cell cycle profiles for wild-type, *FBXW7*^-/-^ and *CCNL1*^*OE*^ cells were generated by flow cytometry. This assay identified a decreased proportion of cells in the G2-M phase in both *FBXW7*^-/-^ and *CCNL1*^*OE*^ cell lines, suggestive of a shortened G2-M phase **(Figure 5C, EV3A)**. Next, using the Eg5 kinesin inhibitor monastrol (Mayer et al., 1999) GFP-tubulin labeled cells were arrested in prometaphase overnight, and released before live-cell imaging to measure the timing of mitosis progression. Consistent with hyperactivity of CCNL1-CDK11 complexes, *FBXW7*^-/-^ and *CCNL1*^*OE*^ cell lines completed cell division with a mean of 295 and 255 minutes respectively while wild-type cells took an average of 350 minutes following monastrol washout (**Figure 5D,E and EV_Movie1-3**) (Renshaw et al., 2019). Importantly, expression level of CCNL1 correlated with mitosis duration (**Figure 5E and 5F**). To further validate this shortened mitosis, the PIP-FUCCI reporter (Grant et al., 2018) **(Figure EV3B)** was employed to assess mitotic timing following nocodazole treatment. Indeed, *FBXW7*^-/-^ and *CCNL1*^*OE*^ cells exited mitosis faster following nocodazole treatment as detected by quantification of mCherry fluorescence (labelling cells in S and G2/M phases) at the bulk population level **(Figure EV3C)**, as well as in individual cells exiting mitosis **(Figure EV3D)**.

**Figure 5.**
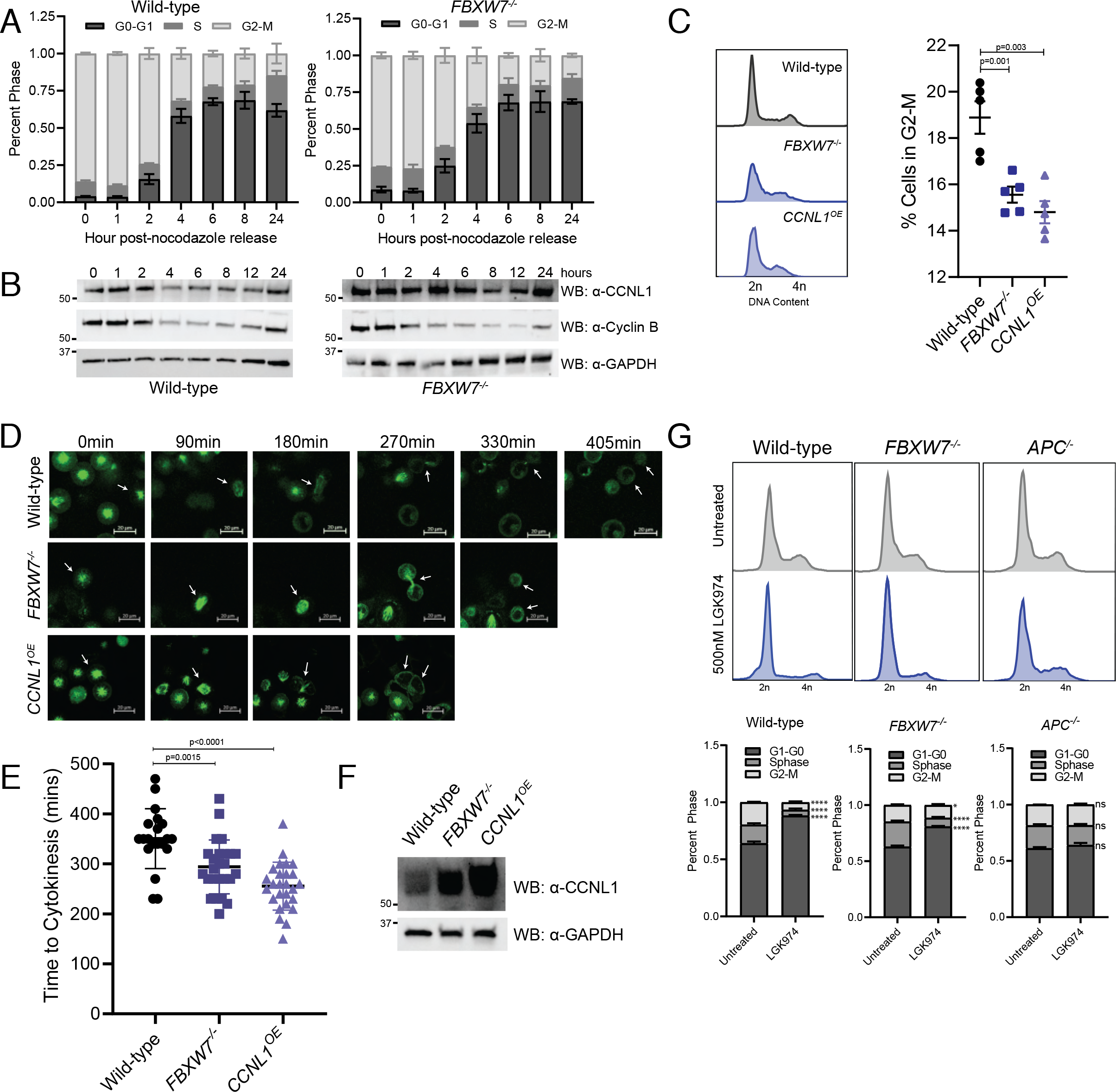
CCNL1 regulates mitotic timing. A Cell cycle profile of nocodazole treated and released HPAF-II wild-type and *FBXW7*^*-/-*^ cells measured by flow cytometry, n=3 independent replicates, mean± SEM. B Immunoblot of HPAF-II wild-type and *FBXW7*^*-/-*^ released from overnight nocodazole treatment at indicated time points, representative of 3 independent replicates C Representative cell cycle profile of HPAF-II wild-type, *FBXW7*^*-/-*^, and *CCNL1*^*OE*^ cells stained with propidium iodide, with quantification of G2-M phase. Three independent replicates, mean ± SEM, one-way ANOVA. D Representative images of GFP-tubulin labeled wild-type, *FBXW7*^*-/-*^, and *CCNL1*^*OE*^ HPAF-II cells monitoring cellular progression through cytokinesis following arrest in prometaphase using monastrol (150μM for 18 hours). Arrows indicate tracked cell, 20μm scale bars E Quantification of cytokinetic timing, n=20, 25, 26 (wild-type, *FBXW7*^*-/*^, *CCNL1*^*OE*^) pooled from three independent experiments, one-way ANOVA. F Immunoblot of wild-type, *FBXW7*^*-/-*^ and *CCNL1*^*OE*^ HPAF-II cell lines demonstrating varying levels of CCNL1. G Cell cycle profiles of wild-type, *FBXW7*^*-/-*^ and *APC*^*-/-*^ HPAF-II cells following treatment with 500nM LGK974 for 24 hours. Representative images of 3 replicates, mean ± SEM, 2-way ANOVA, *p=0.044, ***p<0.001.

Considering a role for CCNL1 in mediating the final stages of mitosis, we wanted to assess the cell cycle profiles of HPAF-II wild-type, *FBXW7*^*-/-*^ and *APC*^*-/-*^ cells following treatment with LGK974, previously reported to arrest cells in G0 (Steinhart et al., 2017). Indeed, HPAF-II *FBXW7*^*-/-*^ cells showed a higher proportion of actively dividing G2-M cells, with an average of 6.5% of G2-M cells in the wild-type and 11% G2-M cells in the *FBXW7*^*-/-*^ cell line following LGK974 treatment, suggesting that cells harboring an *FBXW7*-knockout or LOF mutation may bypass LGK974-induced cell cycle arrest by maintaining a pool of actively dividing cells (**Figure 5G, EV4A**).

### *FBXW7* loss-of-function and *CCNL1* overexpression sensitize cells to CDK11 inhibitor OTS964

A kinase inhibitor currently in preclinical development, OTS964, was recently identified to target CDK11, the cyclin-dependent kinase (CDK) partner of CCNL1 (Lin et al., 2019). Considering the requirement for *CCNL1* in *FBXW7*^-/-^ HPAF-II cells, we aimed to assess whether inhibiting CDK11 using OTS964 could target this synthetic lethal interaction. *FBXW7*^-/-^ HPAF-II cells were isolated and treated with OTS964 in a clonogenic growth assay, and were shown to be hypersensitive to CDK11 inhibition when compared to wild-type cells (**Figure 6A**). Similarly, two independent *CCNL1*^*OE*^ clones demonstrated enhanced sensitivity to OTS964 when compared to wild-type cells, suggesting that CCNL1 expression through *FBXW7* LOF or genetic amplification sensitize cells to CDK11 inhibition (**Figure 6B**).

**Figure 6.**
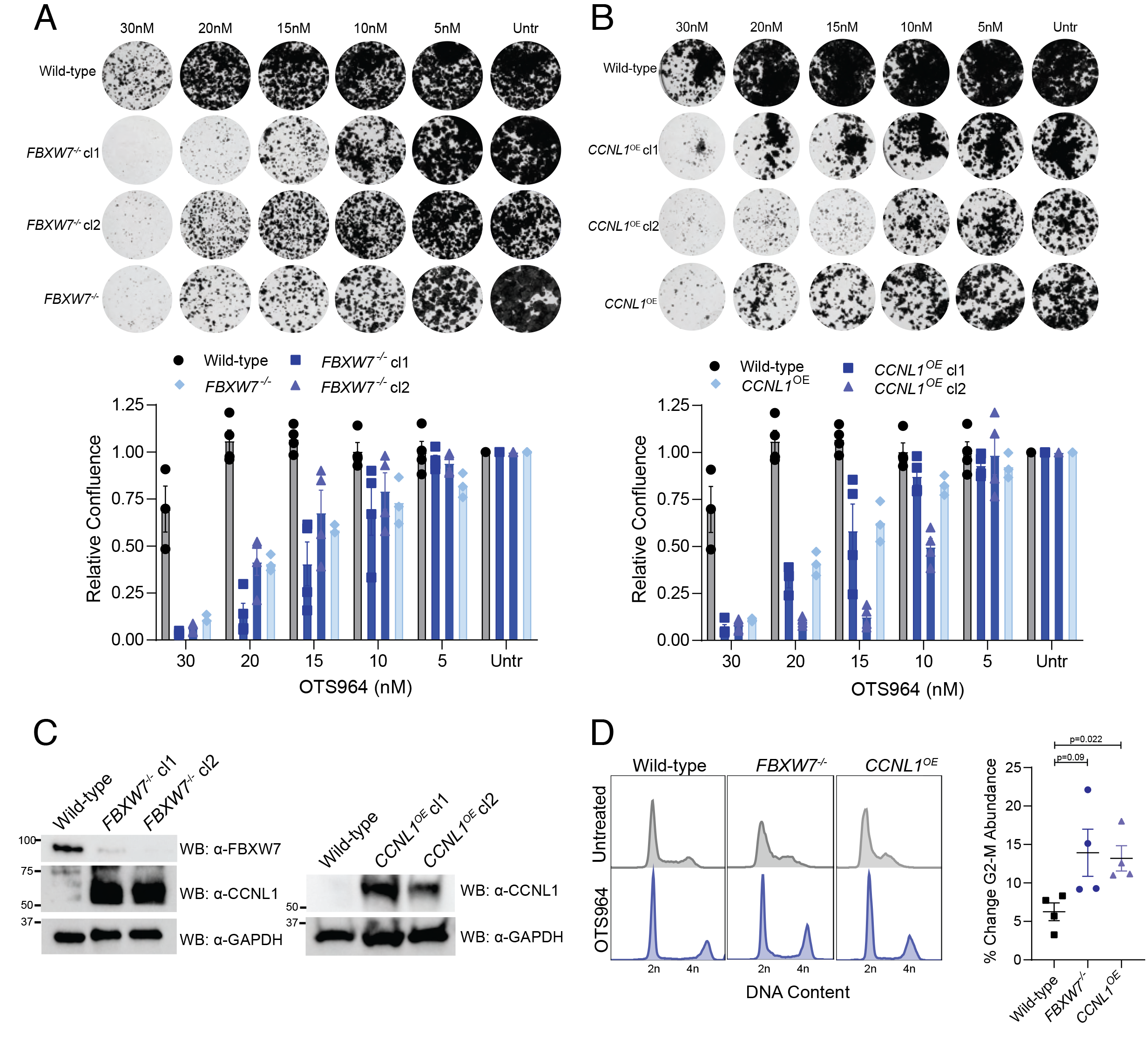
Cells harboring *FBXW7*-LOF mutations or CCNL1 overexpression are highly sensitive to CDK11 inhibitor, OTS964. A Clonogenic growth assays for HPAF-II wild-type and *FBXW7*^*-/-*^ polyclonal, and *FBXW7*^*-/-*^ clones in presence of various OTS964 doses for 14 days. Representative images of 4 independent replicates, quantified by crystal violet absorbance at A595 and plotted mean ± SEM. B Clonogenic growth assays for HPAF-II wild-type and *CCNL1*^*OE*^ polyclonal, and *CCNL1*^*OE*^ clones in presence of various doses of OTS964 for 14 days. Representative images of 4 independent replicates, quantified by crystal violet absorbance at A595 and plotted mean ± SEM. C Immunoblot for the indicated proteins from lysates extracted from the individual *FBXW7*^*-/-*^ and *CCNL1*^*OE*^ clones. D Cell cycle distribution plots with and without 24h OTS964 treatment, representative of 4 independent replicates. Normalized to untreated, mean ± SEM, one-way ANOVA.

To understand the mechanism behind OTS964 sensitivity, cell cycle profiles of wild-type, *FBXW7*^-/-^ and *CCNL1*^*OE*^ cells were obtained by flow cytometry following 24 hours of OTS964 treatment. We identified that OTS964 leads to accumulation of cells in the G2-M phase of the cell cycle, consistent with previous findings (Lin et al., 2019). Interestingly, G2-M accumulation following OTS964 treatment was higher in *FBXW7*^-/-^ and *CCNL1*^*OE*^ cells when compared to wild-type cells suggesting that hyperactivity of CCNL1:CDK11 complexes in G2-M in these genotypes represent a tractable therapeutic vulnerability (**Figure 6D, EV4B**). We conclude that loss of post-translational control of CCNL1 levels is an oncogenic event that can be preferentially targeted by CDK11 inhibitor in cancer cells and that CCNL1 levels could represent a biomarker to stratify patients or predict patient response.

### *FBXW7* mutation and *CCNL1* amplification are mutually exclusive

Considering the high mutation burden of *FBXW7* across a wide range of cancers, and the observation that *CCNL1* amplification occurs in many cancer types, we performed an analysis using cBioPortal to determine the probability of a tumor harboring both *FBXW7* alteration or *CCNL1* amplification. This analysis identified that very few tumors harbor alterations in both *FBXW7* and *CCNL1*, indicating mutual exclusivity and suggesting that these genes function in the same pathway, and that alteration in both would be functionally redundant (**Figure 7A**).

**Figure 7.**
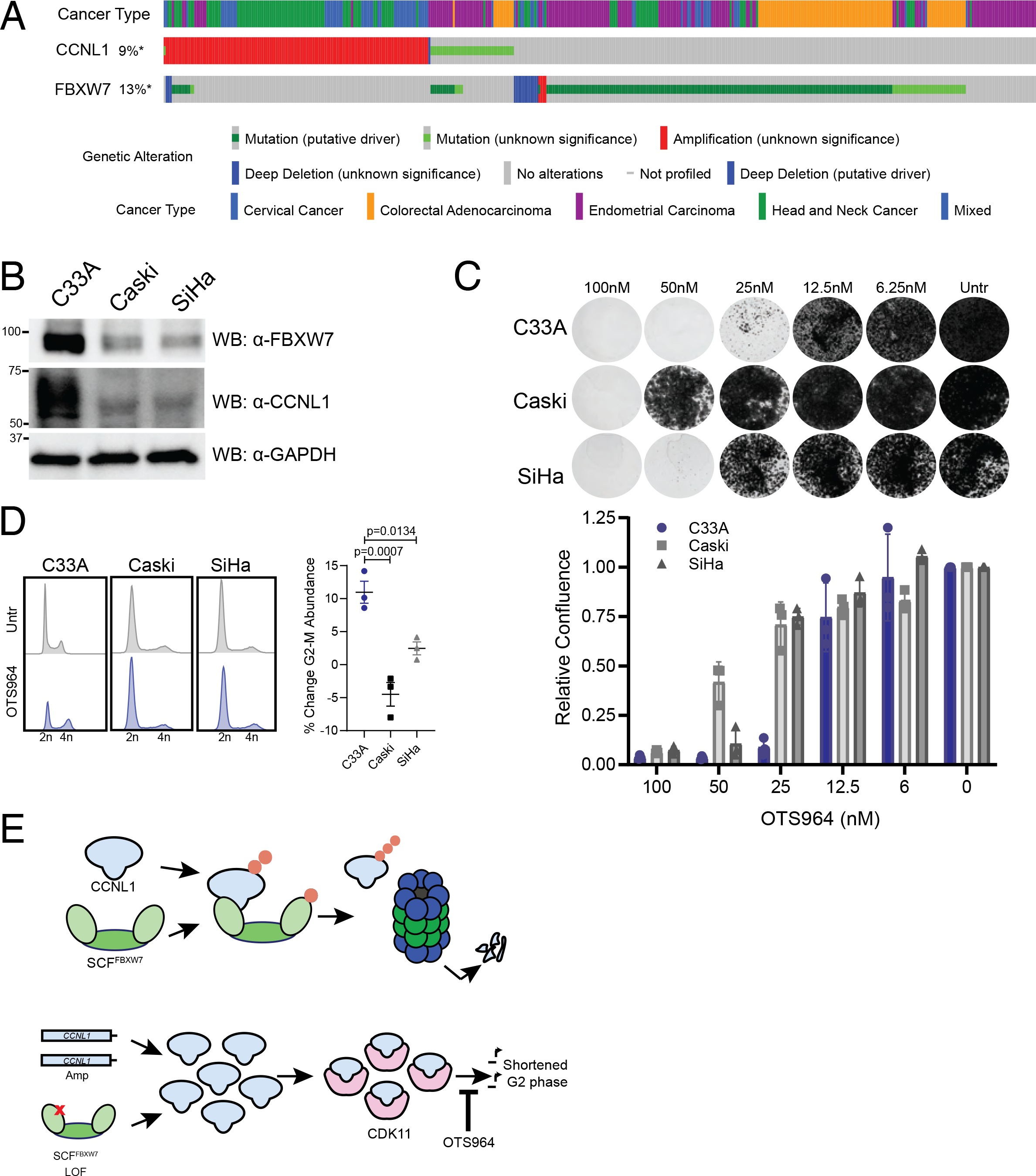
Cervical cancer cell lines exhibit differential sensitivity to OTS964 depending on *FBXW7* mutational status. A Oncoprint from cBioPortal demonstrates mutual exclusivity between *FBXW7* and *CCNL1* alterations B Immunoblot of lysates from cervical cell lines demonstrating C33A cells (*FBXW7*^R465H^) express high levels of CCNL1, representative of three independent replicates. C Clonogenic growth assay of C33A, Caski, and SiHa cells in presence of various doses of OTS964 for 14 days. Representative images of 3 independent replicates, quantified by crystal violet absorbance at A595 and plotted mean ± SEM. D Cell cycle distribution plots with and without 24h OTS964 treatment, representative of 3 independent replicates. Normalized to untreated, mean ± SEM, one-way ANOVA. E Model of proposed mechanism.

### FBXW7-CCNL1-CDK11 axis is therapeutically relevant in cervical cancer cell lines

Given the high prevalence of *FBXW7* mutations and *CCNL1* amplification in cervical cancers (**Figure 7A**), we next screened a genotypically diverse panel of cervical cancer cell lines for their susceptibility to OTS964. The C33A cell line carries a heterozygous *FBXW7*^R465C^ LOF mutation and exhibited high levels of CCNL1 expression when compared to Caski and SiHa, which are wild-type at this locus (**Figure 7B**). Supporting a deregulation of the CCNL1-CDK11 axis, the C33A cell line exhibited increased sensitivity to the CDK11 inhibitor OTS964 than the other *FBXW7*^WT^ lines Caski and SiHa (**Figure 7C**). C33A cells showed an accumulation of cells in the G2-M phase of the cell cycle upon addition of OTS964, whereas Caski and SiHa cell lines show minimal effect (**Figure 7D, EV4C**) - validating the sensitivity of this cell line to OTS964 and perturbation of the G2-M transition. Further, to confirm the on-target toxicity of OTS964 in C33A cells, we performed a genome-wide chemogenomic CRISPR screen using an IC50 dose of OTS964; this screen identified *CDK11A* and *CCNL1* as important in mediating response to OTS964 **(Figure EV4D)**. We conclude that CCNL1 levels could represent a biomarker to predict response to CDK11 inhibitors for treatment of cancers that harbor LOF mutations in *FBXW7* or *CCNL1* amplification **(Figure 7E)**.

## Discussion

The study of the tumor suppressor gene *FBXW7* as substrate specific receptor of a SCF complex has garnered significant interest, and various oncogenic substrates of FBXW7 have been identified since its discovery (Koepp, 2001; Takada et al., 2017; Wei et al., 2005; Welcker et al., 2004a, 2004b). Mechanistically, the substrate phosphodegron CPD motif has also been studied in depth, which describes the I/L-I/L/P-TPXXS motif as the canonical binding moiety required for substrate recognition, in which phosphorylation at the threonine and/or serine residues were described to be important for FBXW7 binding (Nash et al., 2001). The degron motif identified and validated in this work for CCNL1 matches the consensus CPD motif, with the exception that CCNL1 does not contain the I/L-I/L/P sequence directly upstream of the TPXXS sequence. Our work highlighted a requirement for dual phosphorylation at both the threonine and serine residues within the CCNL1 CPD for recognition by FBXW7, similar to what was observed for c-Jun (Wei et al., 2005). This is in contrast to other FBXW7 substrates such as cyclin E (Figure 3E) and c-Myc (Welcker et al., 2004b) where phosphorylation at only one site is sufficient. The functional implications of this differential requirement (single vs dual phosphorylation) is currently unknown but suggests a possible additional layer of regulation through the activity of a priming kinase or other signaling events. Additionally, this data supports the role of FBXW7 in regulating the G2-M transition through control of CCNL1 protein levels - a novel role for this ubiquitous E3 ligase.

Despite early evidence that *CCNL1* functions as an oncogene in a subset of cancers (Muller et al., 2006; Redon et al., 2002; Sticht et al., 2005), relatively little progress has been made towards understanding the molecular mechanisms underlying its role in tumor progression or cell cycle progression. In this study, using an engineered cell model of *FBXW7* LOF and cervical cancer cell lines, harboring wild-type and mutated genotypes for *FBXW7*, we confirmed the mitotic role of CCNL1 (Renshaw et al., 2019) and revealed a molecular basis for deregulation of the FBXW7-CCNL1 axis in cancer. Indeed, both FBXW7 LOF mutations and *CCNL1* amplification would be predicted to result in upregulation of CCNL1 levels and hyperactivation of CDK11. Interestingly, *FBXW7* mutations and *CCNL1* amplification events are mutually exclusive and are highly prevalent in many cancer types. Cervical squamous cell carcinoma appears to be a cancer with frequent alteration in this axis (38% of FBXW7 mutations and 10% of *CCNL1* amplification), which could benefit from CDK11 inhibitor treatment or from other therapies directed at this pathway. Uterine Carcinosarcoma (UCS), an aggressive gynecological cancer with poor prognosis and treatment options, also present with high frequency of *FBXW7* mutations (38%). This, combined with recent studies in engineered mouse models that suggest concurrent loss of function in *FBXW7* and *PTEN*, is a specific driver of UCS tumorigenesis and aggressive tumor behavior (Cuevas et al., 2019) provide further rationale for targeting this molecular perturbation in this disease. *CCNL1* amplification, on the other hand, most frequently occurs in squamous cell cancers such as primarily lung squamous cell carcinoma (19%), cervical squamous cell carcinoma (10%) as well as head and neck squamous cell carcinoma (9%) (Figure 6A). The *CCNL1* gene resides on chromosome 3q25, a region commonly amplified along with *PIK3CA* (Redon et al., 2001) a known human oncogene. Determining whether *CCNL1* amplification is merely a collateral “passenger” event or is required along with *PIK3CA* for cancer initiation or tumor progression remains to be determined. Whether these tumors are more sensitive to CDK11 inhibitors, such as OTS964, in patients that harbor amplification of the 3q25 genomic region also remains to be tested.

Precision oncology is a burgeoning field currently limited by the scarcity of genetic vulnerabilities identified across multiple cancer types. In addition, identification of biomarkers that may predict response to targeted treatments is especially important for stratifying patients for clinical trials to assess the true benefit of a new therapeutic modality. This work has identified a novel synthetic lethal genetic interaction that has the potential to impact a broad range of cancer types. In this study we have demonstrated that CCNL1 expression in cells can predict sensitivity to a novel CDK11 inhibitor, OTS964. Originally identified as a novel inhibitor of the kinase TOPK (T-LAK cell originating protein kinase) (Matsuo et al., 2014), it is unlikely that the activity of OTS964 in our HPAF-II model is based on TOPK targeting, considering this gene is non-essential in both HPAF-II wild-type and *FBXW7*^-/-^ cells and that a chemogenomic CRISPR screen confirmed the on-target activity of OTS964 (**Fig. EV4D**). The importance of CDK11 in cancer progression has been predicted for many years, and our study outlines a novel targeting strategy to guide the use of CDK11 inhibitors.

This work has identified CCNL1 as a novel substrate of tumor suppressor FBXW7, with implications on the growth requirement of cells and tumors harboring *FBXW7* mutations. Through uncontrolled CCNL1 expression, the mitotic phase of the cell cycle is shortened as a result of increased CDK11 activity, which sensitizes cells to its inhibition. CCNL1 is therefore another FBXW7 substrate along with cyclin E, c-myc and c-jun, which have all been linked to cancer. Understanding the individual roles of these substrates and the cellular and cancer contexts where their deregulation contributes to cancer initiation and progression will be important future work needed to realize the potential of various targeted therapies inhibiting these signaling axes.

## Materials and Methods

### Cell Culture & lentivirus production

HPAF-II, HEK293T, SiHa, and C33A cells (ATCC) were cultured in DMEM (Gibco) + 10% FBS (Gibco) and 5% antibiotic & antimycotic (Gibco), Caski (ATCC) cells were cultured in RPMI (Gibco) + 10% FBS and 1% antibiotic & antimycotic, all at 37°C and 5% humidity. Cells were routinely tested for mycoplasma (Lonza), and authenticated by STR profiling at The Center for Applied Genomics at Sickkids Hospital, Toronto. HEK293T cells were seeded to 60% confluence, and the following day transfected with 6μg target plasmid, 6μg pSPAX (Addgene #12260) and 1ug pMD2.G (Addgene #12259) in 60μg polyethylenimine (Sigma-Aldrich) and Opti-MEM (Gibco). 24 hours post-transfection, media was replaced. Lentivirus was harvested 48 hours post-transfection, filtered through a 0.45µm filter, and aliquoted and stored at −80°C prior to use.

### Cell treatments

Cycloheximide chase: following overnight serum starvation, cells were released into 50μg/ml (HEK293T) or 100µg/ml (HPAF-II) cycloheximide (Sigma-Aldrich) for the indicated time points. Clonogenic assays: OTS964 (Selleck Chemicals) was used to treat cells at indicated concentrations for 14 days, with media refreshed every 3-4 days. MG132 (Sigma-Aldrich) was used at 10µM in HEK293T cells for 10 hours, and 1µM in HPAF-II cells for 18 hours. Nocodazole (Cell Signaling Technologies) was used at 150µg/mL for 18h. Monastrol was used at 150µM for 18 hours. Cells were treated with 500nM LGK974 for 24 hours to assess cell cycle dynamics.

### Genome-wide CRISPR screens

#### Positive Selection

HPAF-II cells expressing Cas9 were infected with the Toronto knockout library version 1 (TKOv1) - a pooled sgRNA lentiviral library (Hart et al., 2015) at a multiplicity of infection of 0.3, in the presence of 8μg/ml polybrene (Sigma-Aldrich) for 24 hours. Cells were treated with 2μg/ml puromycin (Life Technologies) for 48 hours. 7 days post-selection, cells were split into treatment groups - one using an LD90 dose of LGK974 at 20nM, and the second a DMSO (Sigma-Aldrich) control; duplicates were included for both treatment arms. LGK974 treatment was harvested at day 28, and DMSO treatment at day 31. Genomic DNA extracted using the QIAmp DNA Blood Maxi Kit (Qiagen). Genomic DNA samples were amplified, and barcoded using i5 and i7 adaptor primers for Illumina next generation sequencing. Barcoded PCRs were sequenced with the Illumina HiSeq2500. Sequenced gRNAs were mapped to the TKOv1 library using MaGECK 0.5.3, and read counts were normalized by total reads per sample before averaging biological replicates and determining gRNA enrichment.

#### Dropout

HPAF-II WT and *FBXW7*^*-/-*^ cells were infected with the Toronto knockout library version 3 (TKOv3) - a pooled sgRNA lentiviral library (Hart et al., 2017) at a multiplicity of infection of 0.3, in the presence of 8μg/ml polybrene (Sigma) for 24 hours. Cells were treated with 2μg/ml puromycin for 48 hours. Following selection, pooled cells were split into three replicates, and passed every 4 days for 24 days, maintaining 18 million cells per replicate. Cell pellets at T=0, 12 and 24 days were collected, and genomic DNA extracted using the QIAmp DNA Blood Maxi Kit (Qiagen). Genomic DNA samples were amplified, and barcoded using i5 and i7 adaptor primers for Illumina next generation sequencing. Barcoded PCRs were sequenced with the Illumina HiSeq2500 with read depths of 200-fold coverage. Sequenced gRNAs were mapped to the TKOv3 library using MaGECK 0.5.3 (Li et al., 2014). Read counts were normalized and fold-change of gRNA distribution compared to T=0 was calculated using the BAGEL package (Hart and Moffat, 2016). BAGEL analysis was performed, and Bayes Factors were compared between HPAF-II wildtype and *FBXW7*^*-/-*^ cells. Z-scores of differential Bayes Factors between wild-type and *FBXW7*^*-/-*^ were calculated.

#### Chemogenomic

C33A cells were infected with the Toronto knockout library version 3 (TKOv3) at a multiplicity of infection of 0.3, in the presence of 8μg/ml polybrene (Sigma) for 24 hours. Cells were treated with 2μg/ml puromycin for 48 hours. Following selection, pooled cells were split into two arms with two replicates per arm. The first arm was treated with DMSO for 16 days, the second arm was treated with 95nM of OTS964 for 16 days. Cell pellets at T=0, 12 and 24 days were collected, and genomic DNA extracted using the QIAmp DNA Blood Maxi Kit (Qiagen). Genomic DNA samples were amplified, and barcoded using i5 and i7 adaptor primers for Illumina next generation sequencing. Barcoded PCRs were sequenced with the Illumina HiSeq2500 with read depths of 200-fold coverage. Sequenced gRNAs were mapped to the TKOv3 library using MaGECK 0.5.3 (Li et al., 2014). gRNAs inducing resistance or synthetic lethal with OTS964 treatment were assessed using the DrugZ algorithm (Colic et al., 2019).

### Generation of *FBXW7* and *APC* mutant cell line

HPAF-II were transfected via electroporation using the Neon system (ThermoFisher Scientific) under the following conditions; 2μg of DNA (pX330 [Addgene # 42230] - sg*FBXW7* or sg*APC*, see sgRNA table) 1150V, 30ms and 2 pulses. Cells recovered for 2 days in full media before the addition of 100nM LGK974 (Cayman Chemicals). The polyclonal cell lines were validated for editing using TIDE (tracking of insertions and deletions) (Brinkman et al., 2014).

### Generation of *CCNL1* overexpressing cell line

HPAF-II wild-type cells were infected with lentivirus carrying a FLAG-CCNL1 cDNA, in the pLenti-puro vector (Addgene #39481). Cells were infected with an ∼0.3 MOI of lentivirus overnight in the presence of 8ug/ml polybrene. The following day, virus-containing media was removed, and cells were selected in 2ug/ml puromycin for 48 hours.

### Generation of GFP-tubulin cell lines

HPAF-II wild-type, *FBXW7*^*-/-*^ and *CCNL1*^OE^ cells were infected with lentivirus carrying tubulin-GFP cDNA, in the pLKO.1 vector (a kind gift from Dr Jason Moffat). Cells were infected with an ∼0.3 MOI of lentivirus overnight in the presence of 8ug/ml polybrene. The following day, virus-containing media was removed, and cells were selected in 2ug/ml puromycin for 48 hours.

### Clone isolation for HPAF-II *FBXW7*^*-/-*^ and *CCNL1*^OE^

HPAF-II *FBXW7-/-* and *CCNL1*OE cells were seeded at 0.5 cells/well in multiple 96-well plates. Single clones were expanded, and tested for *FBXW7*-knockout and CCNL1 expression by western blot. Two clones were chosen and moved forward to clonogenic growth assays.

### Cell competition assay

HPAF-II wildtype and *FBXW7-/-* cells expressing Cas9 were infected with pLentiguide-2A-GFP or pLentiGuide-2A-mCherry-AAVS1 (kind gifts from Dr Daniel Durocher, Lunenfeld-Tanenbaum Research Institute) lentivirus at an MOI of ∼0.3 in the presence of 8μg/ml polybrene. Cells were infected overnight, and treated with 2μg/ml puromycin for 48 hours. Following selection cells were left to recover for 24 hours. Cells transduced with pLentiguide-2A-GFP targeting *AAVS1* or *CCNL1* were mixed 1:1 with pLentiGuide-2A-mCherry-AAVS1 expressing cells, and GFP:mCherry ratios were measured by flow cytometry (Beckman Coulter CytoFLEX) every 4 days for 16 days. Relative fitness was normalized to AAVS1-infected cells.

### Proliferation assay

HPAF-II wildtype and *FBXW7-/-* cells expressing Cas9 were infected with pLentiguide-2A-GFP virus stocks at an MOI of ∼0.3 in the presence of 8μg/ml polybrene. Cells were infected overnight, and treated with 2μg/ml puromycin for 48 hrs. Following selection, fresh media was added, and cells were left for 24 hours to grow. Cells were seeded to ∼2500 cells/well in triplicate in a 96-well plate, and left overnight to attach. Plates were moved to the Incucyte (Sartorius) and confluence was tracked over time. Cell confluence in each line was normalized to AAVS1-infected cells.

### Western blotting

All samples were lysed in 4X Laemmli Sample Buffer (50mM Tris-HCl pH 6.8, 2% SDS, 10% glycerol, 1% β-mercaptoethanol, 12.5mM EDTA, 0.02% bromophenol blue). Lysates were sonicated, boiled, and centrifuged to pellet insoluble material. Approximately 10μg of protein was loaded per sample on a 4-15% SDS-PAGE Stain-Free TGX precast gel (BioRad). Gels were run at 150V for approximately 60 minutes. Gels were transferred to methanol-activated PVDF (BioRad) at 90V for 120 minutes. Membranes were blocked in 5% milk in Tris-buffered Saline (pH 7.4) + 1% Tween-20 (TBS-T) for 1 hour, and incubated with corresponding primary antibodies overnight (see antibody table). The following day, membranes were washed 4 times in TBS-T, and incubated with corresponding secondary antibodies for 1 hour, in 5% milk in TBS-T, at room temperature with agitation. Membranes were washed, and detected using SuperSignal West Pico PLUS chemiluminescent substrate (ThermoFisher) and imaged on the Chemidoc-MP (BioRad).

#### Cytoplasmic fractionation

800,000 cells from each condition were lysed in ice-cold cytoplasmic extraction buffer (10mM HEPES pH8, 1.5mM MgCl2, 10mM NaCl, 0.5mM DTT, 1mM EDTA) and incubated on ice for 15 minutes. NP-40 was added to a final concentration of 0.05%, lysates mixed thoroughly, and the insoluble fraction was collected by centrifugation.

Cytoplasmic fraction was quantified using the Qubit (ThermoFisher) Protein quantification kit, and stored at −80°C until western blotting.

#### Quantification

all western blot quantification was performed by densitometry in ImageJ (FIJI).

### Live Cell Imaging

GFP-tubulin expressing cells were plated into 8-well chamber slides and left overnight to adhere. Cells were then incubated with 150μM monastrol (Selleck Chemicals) overnight. The following day, monastrol was washed out, and cells were imaged every 10 minutes for 8 hours on the Evos FL Auto2 (ThermoFisher) at 20X magnification, at 37°C with 5% CO2. Representative movies were imaged at 37°C and 5% CO2 on a laser scanning confocal microscope (LSM700, Carl Zeiss) at 8-bit with Plan-Apochromat 63X/1.4NA oil immersion objective using Zen software. Z-stacks were captured every 15 minutes for 8 hours. Images were compiled in ImageJ (FIJI).

#### PIP-FUCCI imaging

PIP-FUCCI expressing cells were seeded into 6-well plates. Cells were then treated with 150nM nocodazole overnight. The following day, nocodazole was removed, and cells were imaged every 30 minutes in the Incucyte (Sartorius). mCherry expression was quantified within the incucyte software, and plotted over time.

### Two-dimensional Cell Cycle Flow Cytometry

Cells were grown in logarithmic proliferation, and harvested using 0.025% Trypsin-EDTA. Cells were washed in ice-cold PBS, and fixed in 70% ethanol under vortex, and stored at −20°C overnight. The following day, cells were washed 2x in ice-cold PBS, solubilized in PBS + 1% BSA + 0.15% Triton-X for 15 minutes on ice. Cells were washed and incubated with anti-phosphoH3 (Ser10) (CST) antibody for 1.5h on ice. Cells were washed 2X in PBS + 1% BSA, and incubated with anti-Rb-Alexa488 (ThermoFisher) for 1h on ice. Cells were washed and incubated with 20µg/mL RNAse A (Invitrogen) and 50µg/mL propidium iodide (BioShop) for 30 minutes prior to acquisition on a Beckman Coulter CytoFLEX flow cytometer. Cells were gated for singlets, and cell cycle phase was determined using the intensity in the PE channel. G2-M cells were quantified by gating on all 4N within the pH3+ region.

### Nocodazole Release Cell Cycle Flow Cytometry

Cells were treated with 150µg/ml nocodazole (Cell Signaling Technology) for 18 hours. Following synchronization, cells were released into full medium. Samples were collected for western blotting and flow cytometry at indicated time points. For flow cytometry, samples were trypsinized, collected and washed twice in PBS. Cells were fixed with ice-cold 70% ethanol under vortex, and stored at −20°C. Cells were washed twice in PBS and stained in 50µg/ml propidium iodide (BioShop) in 25nM RNAse A (Invitrogen) in PBS. Samples were run on a Beckman Coulter CytoFLEX flow cytometer. Cells were gated for singlets, and cell cycle phase was determined using the intensity of propidium iodide in the PE channel.

### Immunoprecipitations

Following treatments, 15cm plates were scraped on ice in 1mL PBS and cells collected. Pellets were stored at −80°C until processing. For non-denaturing lysis, pellets were resuspended in RIPA buffer (0.1% SDS, 0.1% NP-40, 2mM EDTA,150mM NaCl, 50mM Tris-HCl pH7.6, 0.5% sodium deoxycholate, 1X protease inhibitor, 10mM NaF, 0.25mM NaOVO_3_), for denaturing lysis, SDS was increased to 1%. Lysates were sonicated, and cleared at 20,000 x g for 20 minutes. Antibodies or FLAG-beads (Sigma-Aldrich) were added to lysates (denaturing lysates first diluted to 0.1% SDS) and incubated at 4°C with end over end rotation for 3 hours. Pre-equilibrated Protein G conjugated agarose beads (Roche) were added for 1 hour.. Beads were collected, washed several times in lysis buffer, and boiled at 95°C for 5 minutes in 4X Laemmli buffer. Samples were stored at −20°C until western blotting.

### Cellular ubiquitination assay

HEK293T cells were transfected with plasmids carrying indicated cDNAs, using PEI. Media was changed the following day. On day 2, cells were starved overnight through the removal of FBS from media. Following overnight starvation, cells were treated with 10µM MG132 (in full media) for 8h. Cells were scraped in ice-cold PBS, and stored at −80°C prior to processing. Lysates were processed as per Immunoprecipitation protocol (denaturing lysis), with the following adjustments: following lysis, samples were boiled at 90°C for 10 minutes, prior to sonication. Elution from Protein-G beads was performed by boiling at 55°C for 5 minutes.

### Fluorescence Polarization Assay

FITC-CCNL1^321-332^ peptides (numbering according to Uniprot Q9UK58-1) were purchased from GenScript and FITC-cyclin E^377-384^ peptides (numbering according to Uniprot P24864-3) were purchased from BioBasic. Experiments were performed by combining 25nM FITC-conjugated peptides and the indicated amount of Skp1-FBXW7^263-707^ complex in buffer containing 25 mM HEPES pH 7.5, 100 mM NaCl, 5 mM DTT, 0.01% Brij-35 and 0.1 mg/mL BSA. Mixed samples (25 μL total volume) were incubated for 30 min in 384-well, black, flat-bottom, low-flange plates (Corning, 3573). Fluorescence intensities were measured using a BioTek Synergy Neo plate reader with excitation and absorbance at 485/528 nm respectively. Fluorescence polarization was calculated with the Gen5 Data Analysis Software. Binding constants for three independent experiments were calculated using GraphPad Prism v8.2.1 (GraphPad) with mean and standard deviation being reported in the figure.

### Statistical tests

All statistical analyses were performed in GraphPad Prism. Data are represented as a mean ± SEM of at least three independent biological replicates.

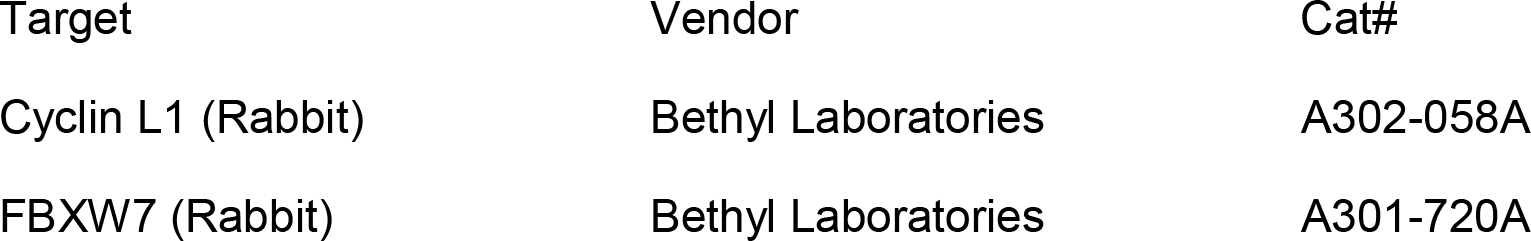
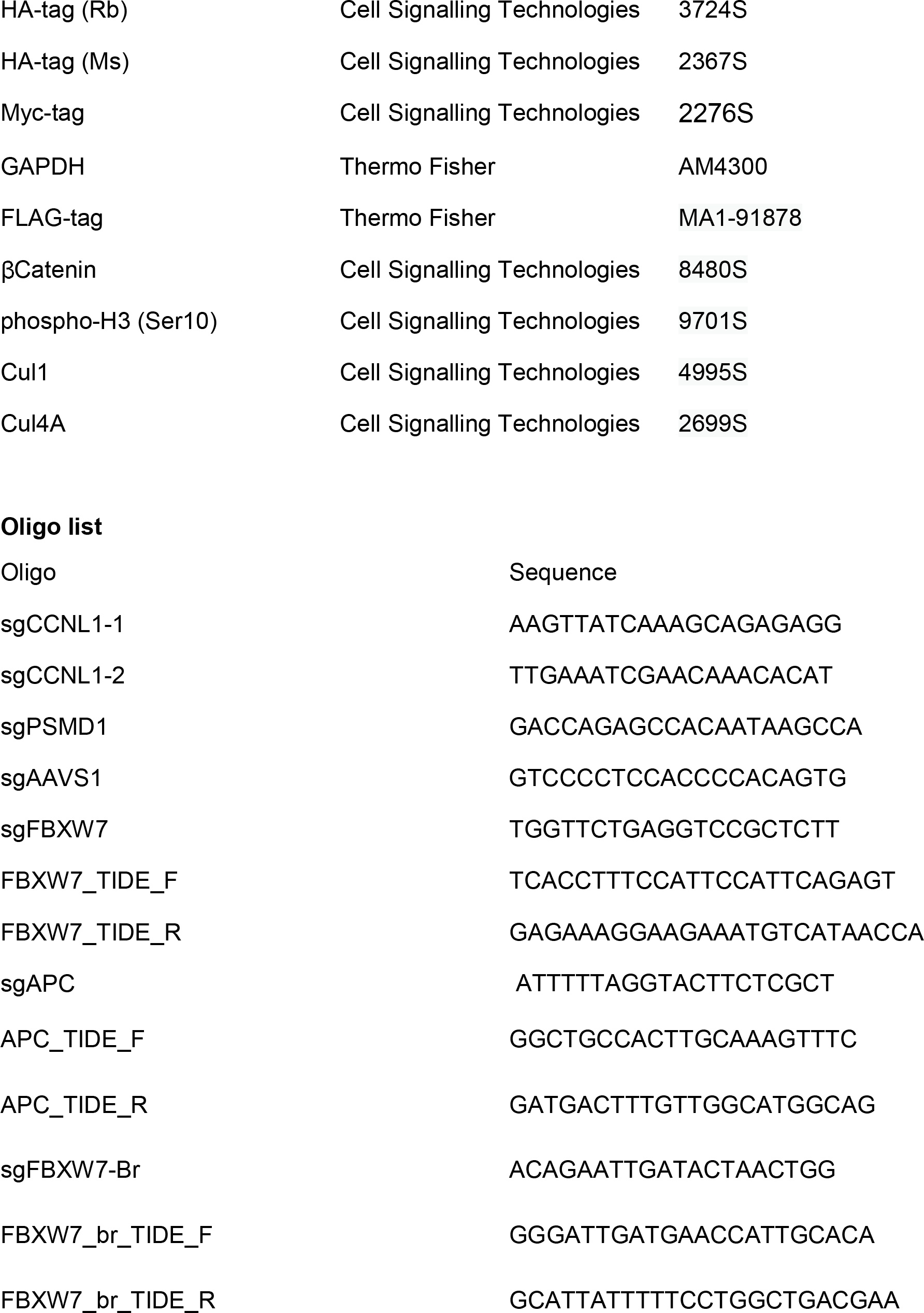

## Supporting information

EV File 1 - Screen Data

## Acknowledgements

This work was supported by grants from the Canadian Institutes of Health Research (PJT-148691 to SA; FDN 143277 to FS), Canadian Cancer Society (CCSRI-Impact grant to FS (704116) and SA (705045) and the Terry Fox Research Institute (to FS). FS is a Canada Research Chair. SO was supported by the Ontario Graduate Scholarship and the Centre for Pharmaceutical Oncology scholarships. Authors would like to thank Andrew Wilde, Michael Ohh, and Brian Raught for reagents, Kin Chan at the LTRI sequencing facility, Azza Al-Mahrouki at the CPO facility, and all members of the Angers lab for support and helpful discussions.

## Author Contributions

Conceptualization: S.O., Z.S., M.M., S.A. Investigation: S.O., S.K., S.Or., Z.S., M.M., S.L., Y.K. Writing - original draft: S.O., S.A. Writing - review & edits: S.O., S.K., M.M., F.S., S.A. Supervision: F.S., S.A.

## Declaration of interests

S.O., Z.S., and S.A. are inventors on a patent involving the use of OTS964 in FBXW7-mutated cancers (WO2021108927A1). F.S. is a founder and consultant at Repare Therapeutics.

## Data Availability

This study includes no data deposited in external repositories. All CRISPR screening data can be found in EV File 1.

## Expanded View

**EV figure 1.**
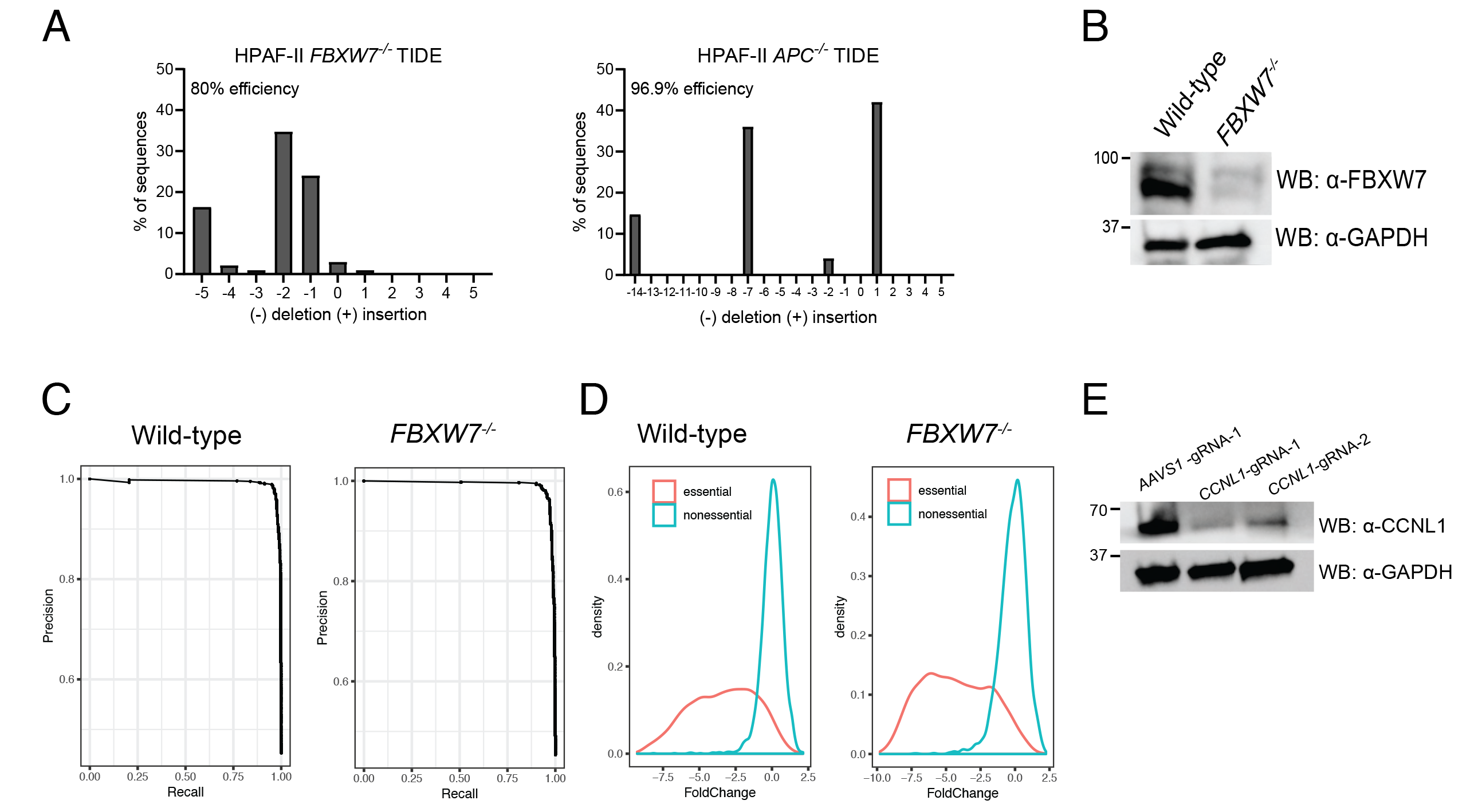
A TIDE analysis of HPAF-II *FBXW7*^*-/-*^ cell line and HPAF-II *APC*^*-/-*^ cell line. B Immunoblot of lysates extracted from HPAF-II *FBXW7*^*-/-*^ cell lines demonstrating knockout of FBXW7 protein expression. C Fold-change plots of HPAF-II wild-type and *FBXW7*^*-/-*^ genome-wide screens demonstrating change in essential genes at T24 of screen. D Precision-recall curves of HPAF-II wild-type and *FBXW7*^*-/-*^ genome-wide screens demonstrating training sets of essential and non-essential genes performed appropriately in the BAGEL algorithm. E Immunoblot of lysates extracted from *FBXW7*^*-/-*^ *c*ells following treatment with sgRNAs targeting *CCNL1*.

**EV figure 2:**
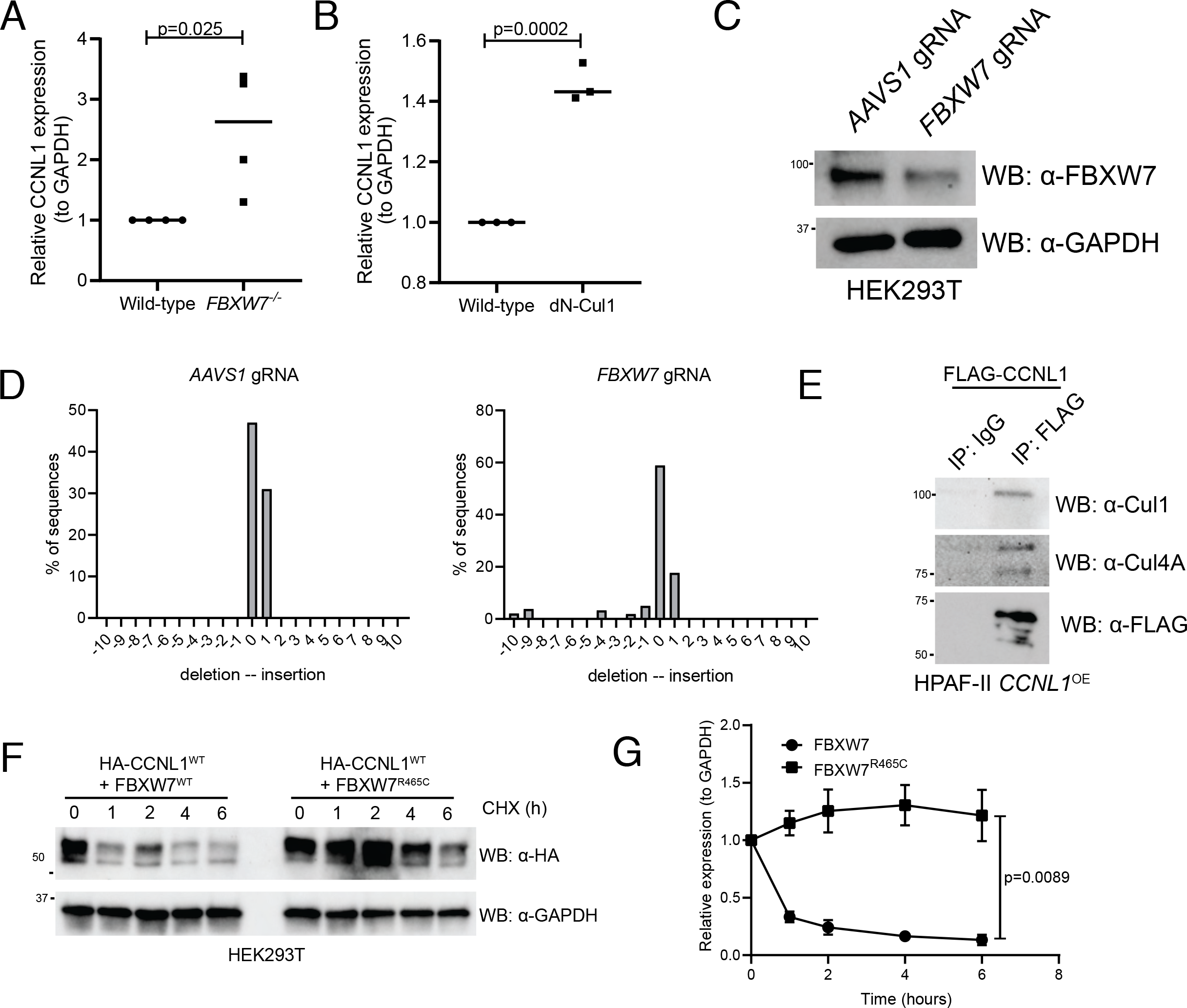
A Quantification of immunoblots in Figure 3A, mean ± SEM, students t-test. B Quantification of immunoblots in Figure 3D, mean ± SEM, students t-test. C Immunoblot analysis of HEK293T cells expressing sgRNAs against *AAVS1* or *FBXW7*. D TIDE analysis of HEK293T cells expressing sgRNAs against *AAVS1* or *FBXW7*. E Immunoblot of immunoprecipitation of FLAG-CCNL1 overexpressed in HPAF-II cells, detecting endogenous Cul1 and Cul4A. Representative image of three independent replicates. F Immunoblot of lysates following cycloheximide treatment of HEK293T cells expressing HA-CCNL1 and FLAG-FBXW7 or FLAG-FBXW7^R465C^. Representative blot of three independent replicates G Quantification of cycloheximide chase in F, mean ± SEM of three independent replicates, t-test at T6.

**EV figure 3:**
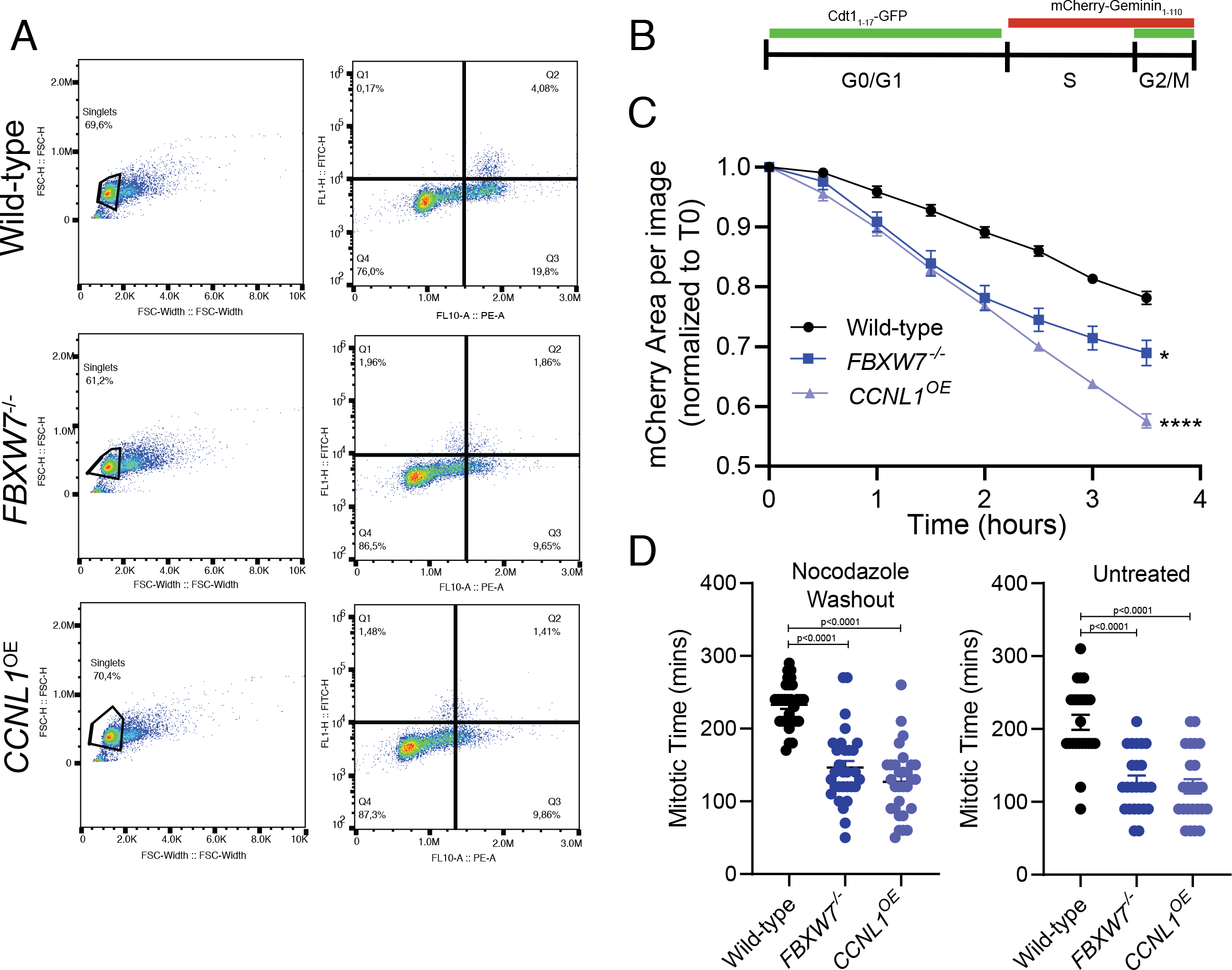
A Representative images of gating strategy for HPAF-II wild-type, *FBXW7*^*-/-*^ and *CCNL1*^*OE*^ cells to determine cell cycle distribution. B Schematic of PIP-FUCCI reporter marker expression through 3 major cell cycle phases. C Live-cell imaging of HPAF-II wild-type, *FBXW7*^*-/-*^ and *CCNL1*^*OE*^ cells expressing PIP-FUCCI reporter treated with nocodazole overnight and released. Images collected over 3.5 hours. Reduction in total population mCherry expression imaged over time, quantified in the Incucyte. Three independent replicates, mean ± SEM, two-way ANOVA. D Live-cell imaging of HPAF-II wild-type, *FBXW7*^*-/-*^ and *CCNL1*^*OE*^ cells expressing PIP-FUCCI reporter either untreated or treated with nocodazole overnight and released. Measurement of individual cells as they lose mCherry expression. n=15 cells per replicate, three independent replicates, one-way ANOVA.

**EV figure 4:**
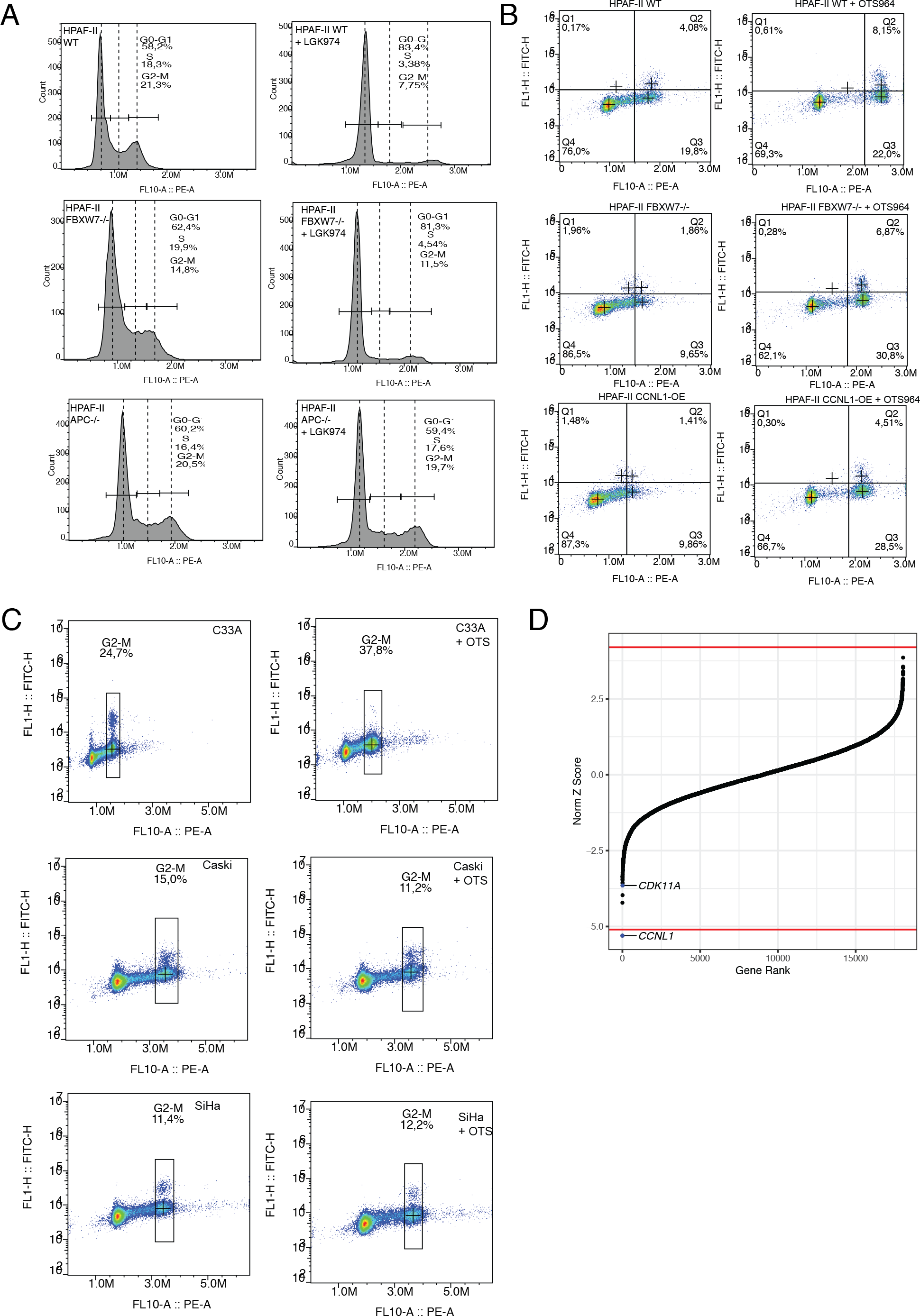
A Representative gating strategy for HPAF-II wild-type, *FBXW7*^*-/-*^ and *APC*^*-/-*^ cells, with and without LGK974 treatment, to determine cell cycle distribution. B Representative gating strategy for HPAF-II wild-type, *FBXW7*^*-/-*^ and *CCNL1*^*OE*^ cells, with and without OTS964 treatment, to determine cell cycle distribution. C Representative images of gating strategy for C33A, Caski and SiHa cells, with and without OTS964 treatment, to determine cell cycle distribution D Normalized Z-score calculated in DrugZ plotted against gene rank for C33A chemogenomic screen with OTS964. Negative score indicates gene knockout synergistic with OTS964, positive scores indicates gene knockouts resistant to OTS964. Red line marks cutoff of FDR<0.05.

**EV Movie 1**. Monastrol wash-out and completion of cytokinesis in GFP-tubulin labelled HPAF-II wild-type cells.

**EV Movie 2**. Monastrol wash-out and completion of cytokinesis in GFP-tubulin labelled HPAF-II *FBXW7*^*-/-*^ cells.

**EV Movie 3**. Monastrol wash-out and completion of cytokinesis in GFP-tubulin labelled HPAF-II *CCNL1*^*OE*^ cells.

**EV File 1:** Raw read counts from HPAF-II LGK974 chemogenomic screen, Bayes Factors from HPAF-II wild-type and *FBXW7*^*-/-*^ genome-wide fitness screens, and raw read counts from C33A cell line OTS964 chemogenomic screen.

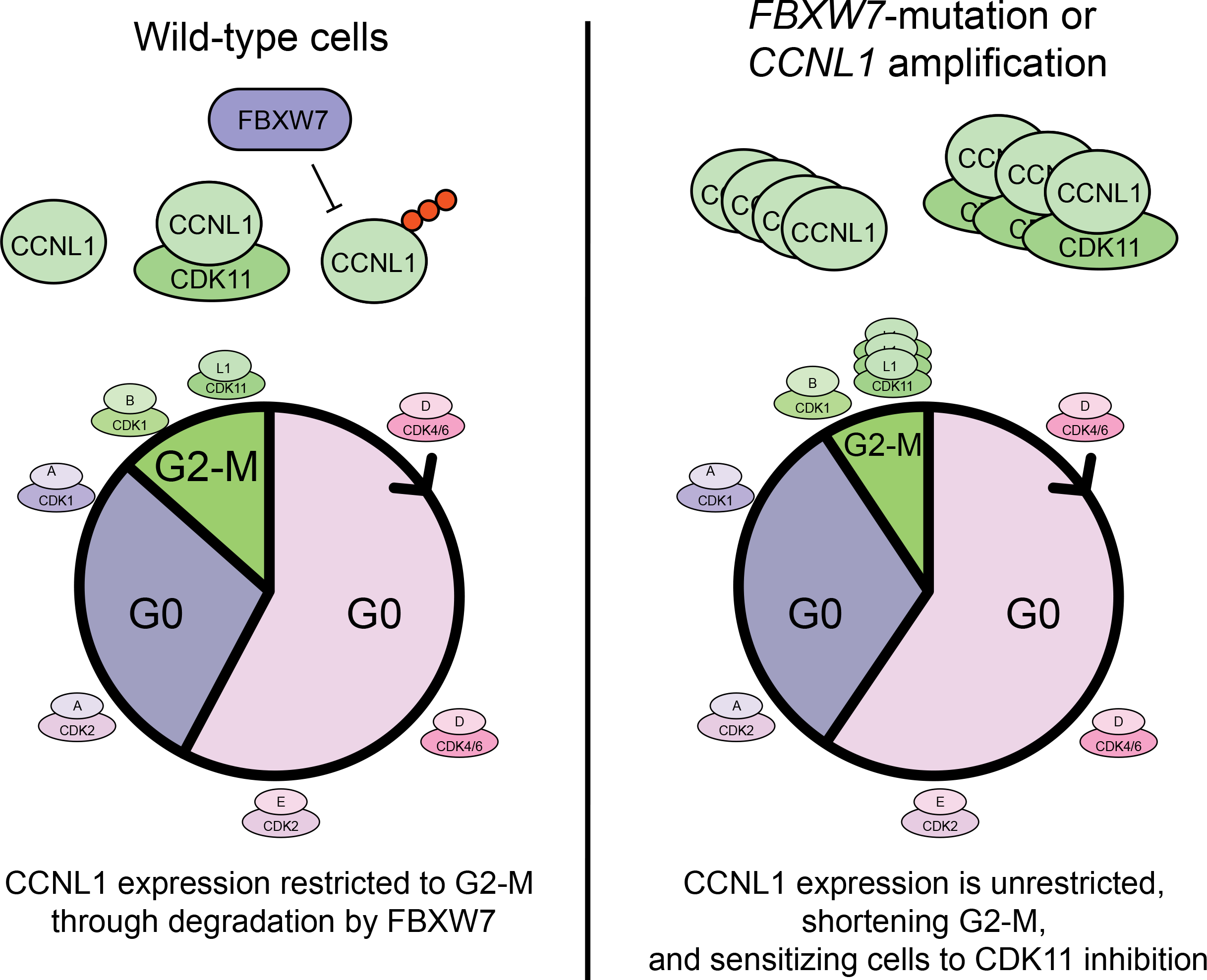

## Notes

### Competing Interest Statement

The authors have declared no competing interest.

